# MARCKS domain phosphorylation regulates the differential interaction of Diacylglycerol Kinase ζ with Rac1, RhoA and Syntrophin

**DOI:** 10.1101/2020.07.08.194100

**Authors:** Ryan Ard, Jean-Christian Maillet, Elias Daher, Michael Phan, Radoslav Zinoviev, Robin J. Parks, Stephen H. Gee

**Author notes:** Department of Biology, University of British Columbia (Okanagan), Kelowna, BC V1V 1V7. Corresponding author: Stephen Gee.

## Abstract

Cells can switch between Rac1, lamellipodia-based and RhoA, blebbing-based migration modes but the molecular mechanisms regulating this choice are not fully understood. Diacylglycerol kinase ζ (DGKζ), which phosphorylates diacylglycerol to yield phosphatidic acid, forms independent complexes with Rac1 and RhoA, selectively dissociating each from RhoGDI. DGKζ catalytic activity is required for Rac1 dissociation but is dispensable for RhoA dissociation. Instead, DGKζ functions as a scaffold that stimulates RhoA release by enhancing RhoGDI phosphorylation by protein kinase Cα (PKCα). Here, PKCα-mediated phosphorylation of the DGKζ MARCKS domain increased DGKζ association with RhoA and decreased its interaction with Rac1. The same modification increased binding of the DGKζ C-terminus to the α1-syntrophin PDZ domain. Expression of a phosphomimetic DGKζ mutant stimulated membrane blebbing in mouse embryonic fibroblasts and C2C12 myoblasts, which was augmented by inhibition of endogenous Rac1. DGKζ expression in differentiated C2 myotubes, which have low endogenous Rac1 levels, also induced substantial membrane blebbing via the Rho-ROCK pathway. These events were independent of DGKζ catalytic activity, but dependent upon a functional C-terminal PDZ-binding motif. Rescue of RhoA activity in DGKζ-null cells required the PDZ-binding motif, suggesting syntrophin interaction is necessary for optimal RhoA activation. Collectively, our results define a switch-like mechanism involving DGKζ phosphorylation by PKCα that favours RhoA-driven blebbing over Rac1-driven lamellipodia formation and macropinocytosis. These findings provide a mechanistic basis for the effect of PKCα signaling on Rho GTPase activity and suggest PKCα activity plays a role in the interconversion between Rac1 and RhoA signaling that underlies different migration modes.

## Introduction

Rho GTPases are molecular switches that control a wide variety of signal transduction pathways in eukaryotic cells. They are best known for their pivotal role in regulating the actin cytoskeleton, but they also influence cell polarity, microtubule dynamics, membrane transport pathways, and cell cycle progression (1). These biological functions of the Rho proteins are critically important during tissue morphogenesis events required for the normal development of multicellular organisms and have decisive roles in the invasion and metastasis of cancer cells (2). In cultured mammalian cells Rac1 promotes actin polymerization and focal complex assembly leading to lamellipodia protrusion and membrane ruffle formation, while RhoA promotes the assembly of actin stress fibers and focal adhesions (3,4) and drives actomyosin-based membrane blebbing and microvesicle formation (5).

Rho GTPases cycle between inactive, GDP-bound and active, GTP-bound conformations. The active forms interact with specific downstream effectors to elicit distinct biological responses. Rho GTPase activity is tightly regulated by guanine nucleotide exchange factors (GEFs), which activate Rho proteins by promoting the exchange of GDP for GTP; by GTPase-activating proteins (GAPs), which inactivate Rho proteins by enhancing their intrinsic GTPase activity; and by guanine nucleotide dissociation inhibitors (GDIs), which sequester Rho proteins as soluble cytosolic complexes and prevent the association of their C-terminal lipid moieties with the plasma membrane (6,7).

Our previous studies demonstrate that diacylglycerol kinase ζ (DGKζ), which phosphorylates diacylglycerol (DAG) to yield phosphatidic acid (PA), forms independent signaling complexes with both Rac1 and RhoA and plays a central role in their activation by dissociating them from their common inhibitor, RhoGDI (8,9). We first showed DGKζ is a key component of a signaling complex that includes Rac1, RhoGDI, and the serine/threonine kinase PAK1, which together, function as a Rac1-selective, RhoGDI dissociation factor^21^. In response to growth factor stimulation, DGKζ stimulates the conversion of DAG into PA, which stimulates PAK1 activity (8,10). Active PAK1 phosphorylates RhoGDI on Ser-101 and Ser-174 to trigger Rac1 dissociation, enabling its subsequent activation by plasma membrane GEFs (8,11). A catalytically inactive DGKζ mutant was unable to rescue the decrease in Rac1 activity in DGKζ-null mouse embryonic fibroblasts (MEFs), consistent with the requirement of DGKζ enzymatic activity for Rac1 activation via this mechanism.

DGKζ is also a component of a distinct signaling complex that includes RhoA, RhoGDI, and the serine/threonine kinase protein kinase Cα (PKCα) that functions as a RhoA-selective, RhoGDI dissociation factor (9). RhoA release is mediated by PKCα phosphorylation on RhoGDI Ser-34, which uses a non-canonical method of PKCα activation stimulated by uncleaved phosphatidylinositol 4,5-bisphosphate (PI[4,5])P_2_) (12). Our findings indicate that optimal RhoA activation and function in MEFs requires DGKζ. However, in contrast to Rac1 regulation, DGKζ catalytic activity is dispensable for RhoA-RhoGDI dissociation, suggesting it functions primarily as a scaffold to enhance RhoGDI phosphorylation by PKCα (9). The molecular determinants that mediate selective binding of DGKζ to either Rac1 or RhoA are unknown.

DGKζ and PKCα exist in a regulated signaling complex, wherein DGKζ inhibits PKCα activity by metabolizing DAG, a cognate PKCα activator (13). DGKζ contains a motif similar to the phosphorylation-site domain of the myristoylated alanine-rich C-kinase substrate (MARCKS) protein (14), a Ser/Thr-rich region phosphorylated by PKCα (15). PKCα-mediated phosphorylation of this motif in DGKζ abolishes their interaction and impairs PKCα regulation, allowing unfettered PKCα activity (13). The MARCKS domain in DGKζ is also a bipartite nuclear localization signal; its phosphorylation negatively regulates DGKζ nuclear localization (15). Phosphorylation of this motif also enhances the translocation of cytoplasmic DGKζ to the plasma membrane where its substrate DAG is available (16,17). Despite elevated plasma membrane localization, MARCKS domain phosphorylation reduces DGKζ enzymatic activity by ~50% (18). Thus, PKCα-mediated phosphorylation of the MARCKS domain has pleiotropic effects on DGKζ function.

Since DGKζ is common to both Rac1 and RhoA dissociation mechanisms, we surmised that signals regulating DGKζ activity help to control the balance of Rac1 and Rho activity. Here, we investigated the impact of DGKζ MARCKS domain phosphorylation on the selective regulation of Rac1 and RhoA signaling. We demonstrate that PKCα-mediated phosphorylation of the MARCKS domain increases the interaction of DGKζ with RhoA and with the PDZ domain of α1-syntrophin, while simultaneously decreasing its interaction with Rac1. A DGKζ mutant that mimics MARCKS domain phosphorylation, in conjunction with reduced Rac1 activity, preferentially activated RhoA-driven membrane blebbing, which was dependent upon the DGKζ C-terminal PDZ-binding motif that mediates association with syntrophin. Collectively, these findings reveal a mechanism for the selective binding of DGKζ to Rac1 or RhoA and suggest MARCKS domain phosphorylation functions as an intramolecular switch that triggers conformational changes that activates RhoA over Rac1.

## Results

We first examined whether PKC activity affects the interaction of DGKζ with the Rho GTPases RhoA and Rac1. To specifically assess the impact of PKC activity on the DGKζ/RhoA interaction, we monitored the binding of exogenous, HA-tagged wild-type DGKζ from lysates of mouse embryonic fibroblasts (MEFs) to recombinant glutathione S-transferase (GST) fusion proteins of constitutively active (RhoA^V14^) and inactive (RhoA^N19^) versions of RhoA. Treatment with phorbol myristate acetate (PMA), a potent PKC activator, for 30 min prior to harvesting increased HA-DGKζ binding to GST-RhoA^V14^ and GST-RhoA^N19^ by approximately 2-fold compared to vehicle control (**Fig. 1A-D**). The increase in binding was blocked by the specific PKCα/β inhibitor Gö6976 indicating that PKC catalytic activity mediates this effect. Under the same conditions, PMA treatment caused GST-Rac1^V12^ to capture substantially less HA-DGKζ than controls (**Fig. 1E, F**), consistent with our previous findings (19). Again, this effect was blocked by PKCα/β inhibition. These data suggest PKCα/β-mediated phosphorylation simultaneously increases DGKζ binding to RhoA while decreasing binding to Rac1.

**Figure 1.**
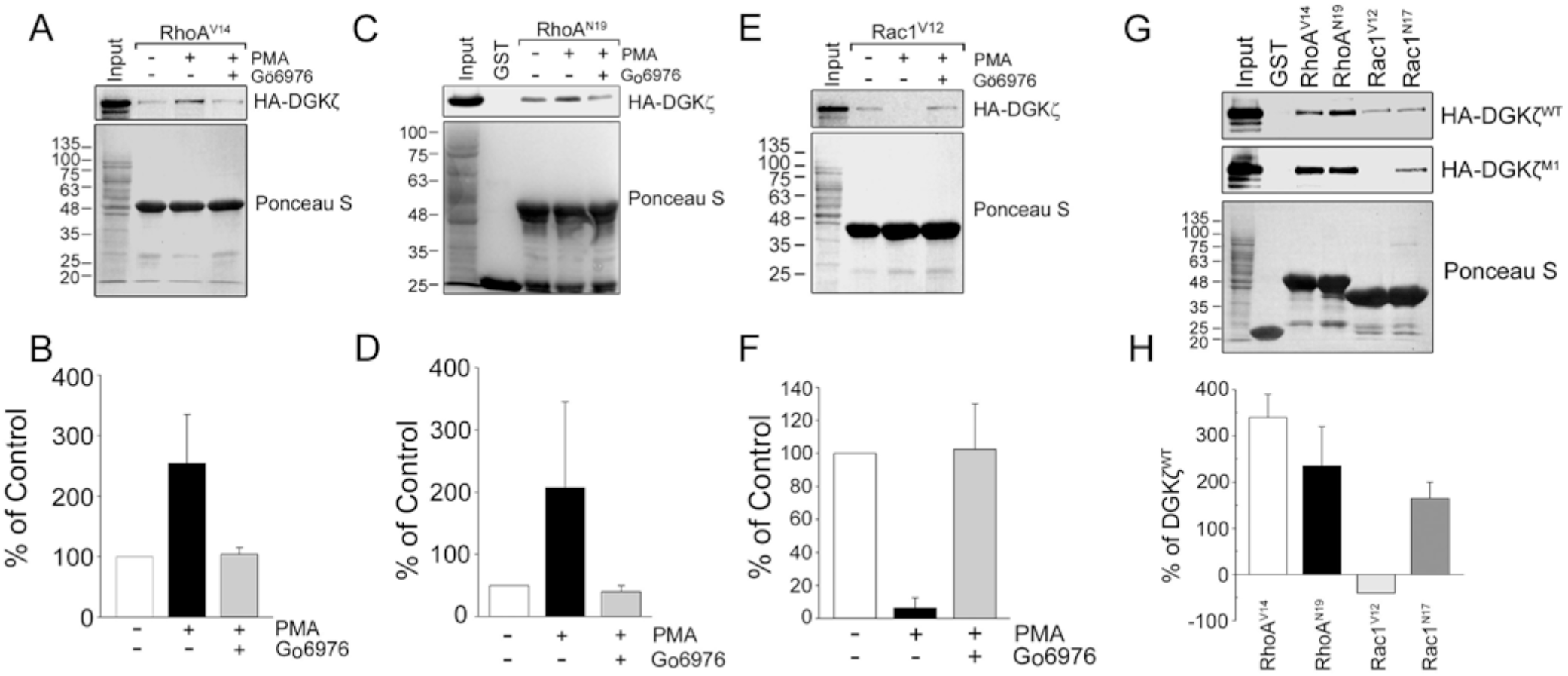
PKCα/β activity differentially modulates DGKζ interaction with Rac1 and RhoA. Wild type MEFs infected with HA-DGKζ were stimulated with 100 ηM PMA or DMSO for 30 min prior to harvesting. The cell extracts were incubated with beads conjugated to (**A**) GST-RhoA^V14^, (**C**) GST-RhoN^19^ or (**E**) GST-Rac1^V12^. Bound DGKζ was analyzed by blotting with anti-HA antibodies. Ponceau S stained blots show the relative amounts of each fusion protein loaded. To inhibit PKCα/β activity, cells were treated with 1 μm Gö6976 for 10 min before PMA stimulation. (**B-F**) Graphs showing the quantification of HA-DGKζ captured by (**B**) RhoA^V14^, (**D**) RhoN^19^ or(**F**) Rac1^V12^ by densitometric analysis of immunoblots. Values are the average percent change compared to unstimulated cells from three independent experiments. Errors bars represent S.E.M. (**G**) Extracts of MEFs infected with HA-DGKζ or HA-DGKζ^M1^ were incubated with beads conjugated to GST or the indicated GST-Rho GTPase constructs. Bound proteins were analyzed as above. (**H**) Graph showing the quantification of captured HA-DGKζ^WT^ or HA-DGKζ^M1^. Values are the average percent change relative to HA-DGKζ^WT^ from a minimum of three independent experiments. Errors bars represent S.E.M.

We showed previously that a DGKζ mutant (DGKζ^M1^) in which all four serine residues in the MARCKS domain were changed to aspartate to mimic phosphorylation (15);(17), bound less efficiently to Rac1^V12^ than wild type DGKζ (19). Here, we compared the binding of HA-tagged wild type DGKζ and DGKζ^M1^ to RhoA. GST-RhoA^V14^ and, to a lesser extent, GST-RhoA^N19^, captured substantially more HA-DGKζ^M1^ than wild type HA-DGKζ, suggesting MARCKS domain phosphorylation increases the interaction of DGKζ with RhoA (**Fig. 1G, H**). In the same experiment, GST-Rac1^V12^ captured wild type DGKζ but no detectable DGKζ^M1^, while GST-Rac1^N17^ captured approximately equivalent amounts of both proteins. Neither protein was captured by GST alone, suggesting the interactions between DGKζ and these Rho GTPases are specific. Collectively, these findings suggest PKCα/β-mediated phosphorylation of the MARCKS domain switches the binding preference of DGKζ from Rac1 to RhoA.

### Syntrophin Interaction

DGKζ contains a C-terminal PDZ-binding motif that mediates its interaction with the syntrophin family of PDZ domain-containing scaffold proteins (20). To investigate whether PKC activity affects their interaction, cells infected with HA-DGKζ^WT^ were treated with vehicle or PMA. The detergent-solubilized cell lysates were incubated with a GST fusion protein of the α1-syntrophin PDZ domain (GST-α1-PDZ) and the amount of bound DGKζ was detected by immunoblotting with an anti-HA antibody. HA-DGKζ binding to GST-α1-PDZ was significantly increased following PMA stimulation (**Fig. A and B**). Applying Gö6976 prior to PMA stimulation reduced the interaction, demonstrating that PKCα/β activity is required for this effect. These results suggest PKCα/β-dependent activity positively regulates the interaction of DGKζ with the α1-syntrophin PDZ domain.

To determine if MARCKS domain phosphorylation by PKC specifically accounts for the observed increase in DGKζ binding to α1-PDZ following PMA stimulation, we compared the binding of α1-PDZ to DGKζ^WT^ and DGKζ^M1^. Lysates of transiently transfected wild type MEFs were incubated with GST alone or GST-α1-PDZ and bound DGKζ was detected and quantified as above. Substantially more DGKζ^M1^ (~2.5-fold) bound to α1-PDZ than did DGKζ^WT^, despite equivalent levels of expression (**Fig. 2C and D**). These data suggest MARCKS domain-phosphorylation accounts for the PMA-induced increase in binding to the α1-PDZ domain.

**Figure 2.**
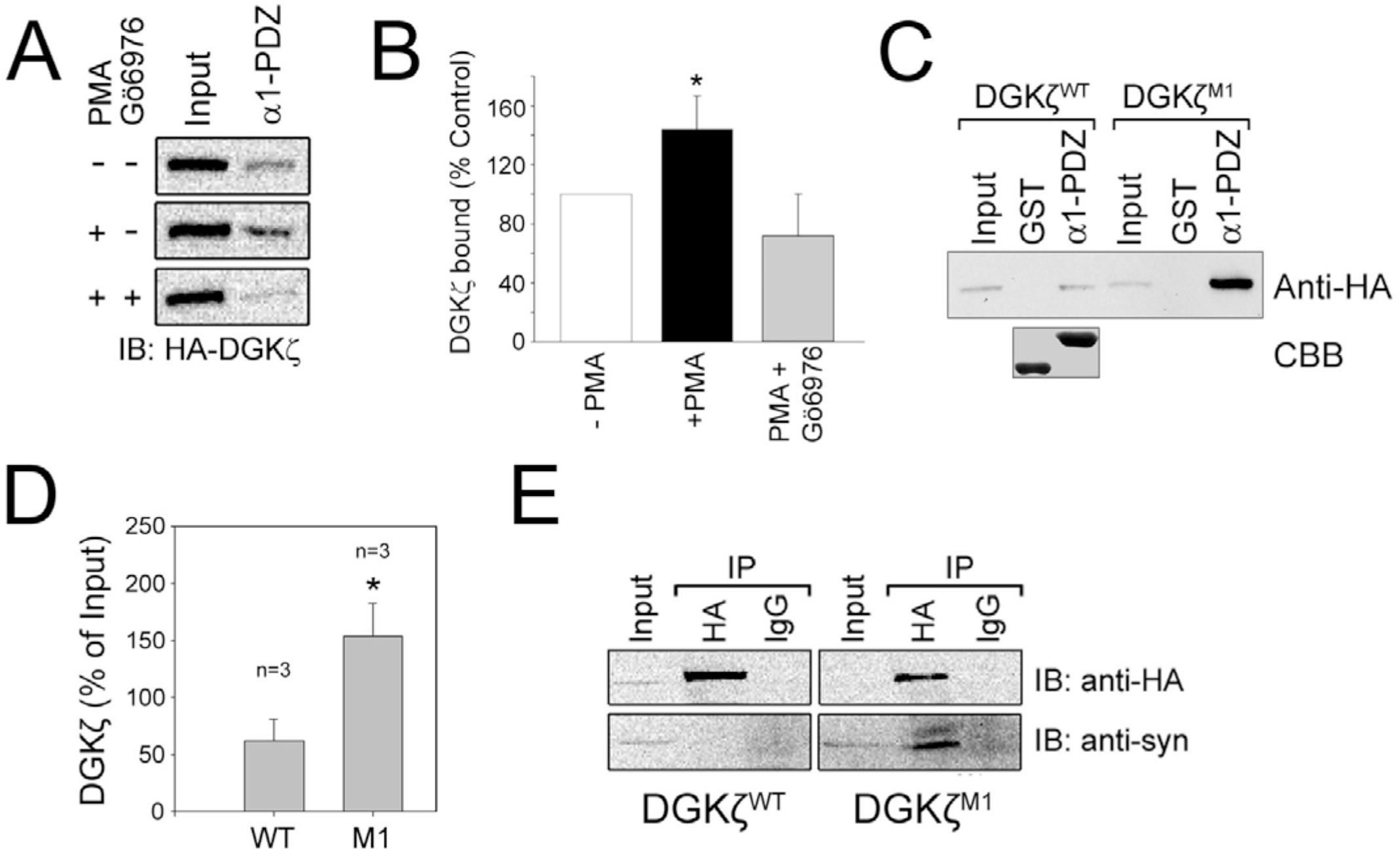
PKC activation increases the association of DGKζ with syntrophin via PDZ interactions. (**A**) Pulldown of HA-DGKζ with the α1-syntrophoin PDZ domain. MEFs infected with an adenoviral vector encoding HA-DGKζ were treated with PMA, with or without Gö6976 for 30 min before lysing. The cell extracts were incubated with GST alone or with GST-α1-PDZ. Bound proteins were detected by anti-HA antibody. Input represents 2% of the starting material. (**B**) Graph showing the amount of bound DGKζ relative to input. Values are the average of three independent experiments. Errors bars represent S.E.M. An asterisk indicates a statistically significant difference from the unstimulated sample (*P* < 0.05, two-tailed *t* test). (**C**) The MARCKS phosphorylation mutant shows increased binding to α1-syn PDZ. Beads charged with GST or GST-α1-PDZ were incubated with extracts of MEF transiently transfected with HA-DGKζ or HA-DGKζ^M1^. Bound proteins were analyzed as above. Input represents 2% of the starting material. Coomassie Brilliant Blue (CBB) staining shows the relative amounts of fusion proteins loaded. (**D**) Graph showing bound DGKζ relative to input. Values are the average of three independent experiments. An asterisk indicates a statistically significant difference from the wild type DGKζ (*P* < 0.05, two-tailed *t* test). (**E**) Increased interaction of endogenous syntrophins with HA-DGKζ^M1^. Extracts of cells infected with HA-DGKζ^WT^ or HA-DGKζ ^M1^ incubated with an anti-HA antibody or control IgG. The immunoprecipitates were analyzed for syntrophin and HA-DGKζ. Input represents 2% of the starting material.

To confirm that MARCKS domain phosphorylation affects the interaction of DGKζ with full-length, endogenous syntrophins, lysates of cells infected with either HA-DGKζ^WT^ or HA-DGKζ^M1^ were immunoprecipitated with an anti-HA antibody and the immune complexes analyzed by immunoblotting with a pan specific syntrophin monoclonal antibody (21). Under these conditions, syntrophins coimmunoprecipitated with HA-DGKζ^M1^ but not with HA-DGKζ^WT^, despite the fact that roughly equivalent levels were immunoprecipitated by the anti-HA antibody (**Fig. 2E**). Syntrophins were not precipitated by control IgG suggesting the interaction is specific. These results suggest MARCKS domain phosphorylation promotes DGKζ interaction with syntrophin. Taken together, these findings reveal MARCKS domain phosphorylation is an important regulatory switch that favors the interaction of DGKζ with RhoA and syntrophin.

### Membrane Blebbing

Since DGKζ^M1^ preferentially interacts with RhoA, we tested if its exogenous expression in MEFs would activate RhoA-ROCK signaling leading to membrane blebbing, a downstream effect of RhoA activity. The majority of cells expressing HA-DGKζ^M1^ were well spread, consistent with Rac1, and not RhoA, activation, however approximately 20% of the cells had blebs, roughly twice as many as in the uninfected MEF control (**Fig. 3A and B)**. Since Rac1 activity suppresses RhoA signaling (22), we hypothesized that DGKζ^M1^ expression and inhibition of Rac1 activity might lead to a further increase in blebbing. Indeed, Rac1 inhibition with 100 μM NSC 23766 in DGKζ^M1^-expressing cells substantially increased the percentage of those with blebs **(Fig. 3A and B**). When MEFs were infected with an adenovirus encoding a constitutively active RhoA mutant, RhoA^V14^, approximately 60% of cells had membrane blebs. The percentage of MEFs infected with the dominant-negative RhoA mutant, RhoA^N17^, that underwent blebbing was not significantly different from uninfected control cells (**Fig. 3B**). Similar results were also obtained in C2C12 mouse myoblasts, in which treatment of cells expressing either DGKζ^WT^ or DGKζ^M1^ with NSC 23766 led to an increase in the percentage of cells with blebs (**Fig. 3C**).

**Figure 3.**
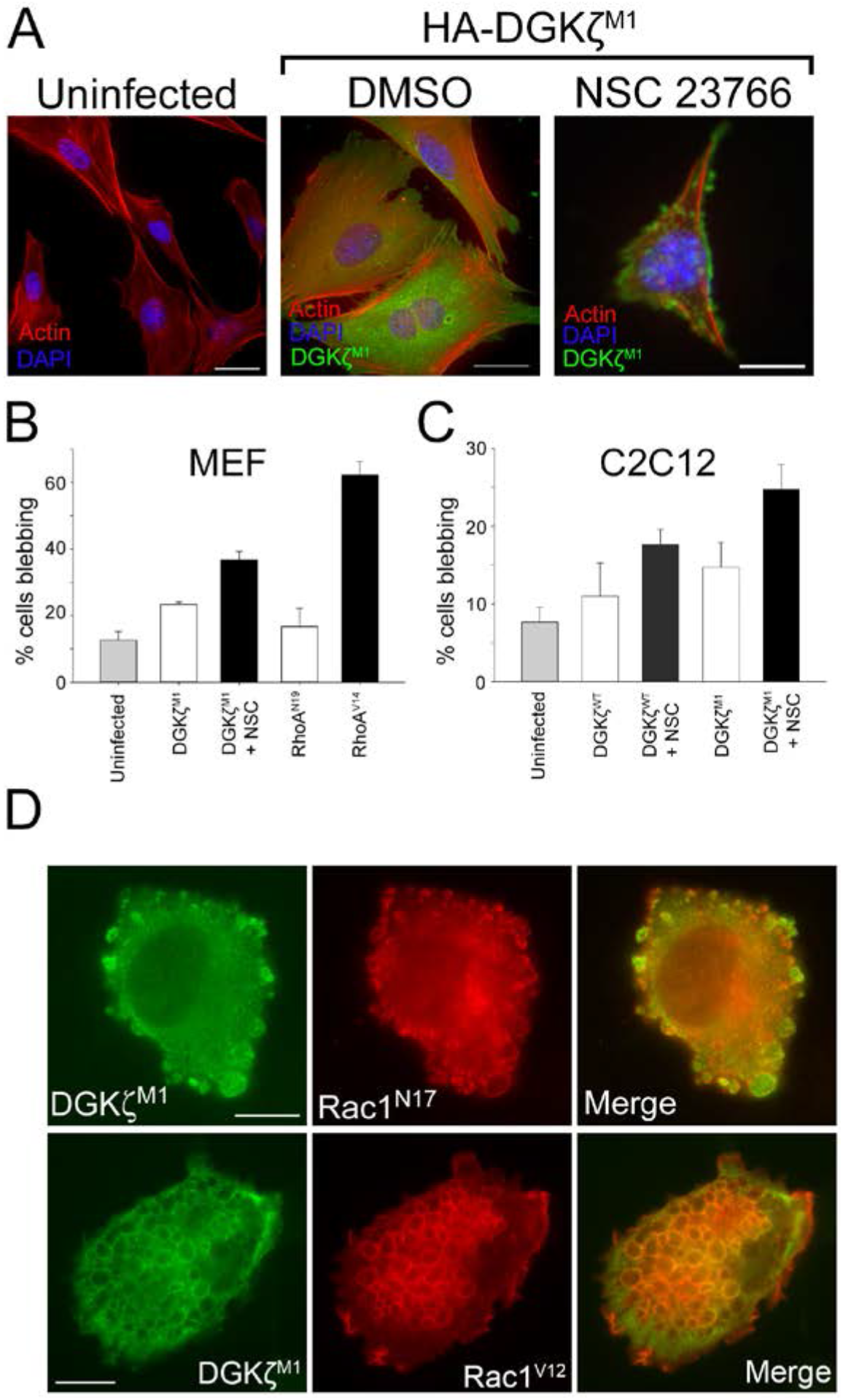
Membrane blebbing in MEFs and C2C12 myoblasts. (**A**) Representative immunofluorescence images of uninfected MEFs or MEFs expressing HA-DGKζ^Μ1^ and treated with vehicle or the Rac1 inhibitor NSC 23766 for an additional 18 hours. The cells were fixed and stained with DAPI (blue), AlexaFluor 594-conjugated phalloidin (red) and anti-HA (green). Scale bars = 20 μm. (**B**) Graph showing the quantification of membrane blebbing in MEFs infected with the indicated adenoviral constructs. Values are the average of two independent experiments. Errors bars represent S.E.M. A minimum of 150 cells were counted for each condition. (**C**) Blebbing quantification in C2C12 infected with the indicated adenoviral constructs, with or without NSC 23766. Values are the average of three independent experiments. Errors bars represent S.E.M. A minimum of 150 cells were counted for each condition. (**D**) Representative images of C2 myoblasts expressing HA-DGKζ^M1^ (green) and myc-Rac1^N17^ or myc-Rac1^V12^ (red). Merged images are shown at the right. Scale bars = 10 μm.

To bolster the idea that Rac1 inactivation leads to increased blebbing, C2 myoblasts were cotransfected with HA-DGKζ^M1^ and a myc-tagged inactive Rac1 mutant, Rac1^N17^, which functions as a dominant-negative by sequestering available GEFs, preventing activation of endogenous Rac1 (23). C2 cells co-expressing HA-DGKζ^M1^ and myc-Rac1^N17^ had many, large membrane blebs. Rac1^N17^ and DGKζ^M1^ were colocalized on bleb membranes and DGKζ^M1^ was additionally found in the bleb cytosol (**Fig. 3D, *top panels***). In contrast, co-expression of DGKζ^M1^ with a constitutively active Rac1 mutant, Rac1^V12^, induced the formation of large macropinosomes consistent with our previous studies (24). Although DGKζ^M1^ has reduced kinase activity and preferentially associates with RhoA, taken together, our finding suggest it continues to activate sufficient levels of Rac1 to promote macropinocytosis. However, when Rac1 is inactive or unavailable, DGKζ^M1^ drives membrane blebbing.

Rac1 activity is high in proliferating C2 myoblasts and decreases during differentiation and fusion of myoblasts into multinucleated muscle fibers (25). We took advantage of this natural occurring change in Rac1 levels to study the effect of exogenous DGKζ expression, driven by adenoviral infection, on membrane blebbing. An adenovirus bearing green fluorescent protein (GFP), used as a control, only induced minimal blebbing in infected C2 myotubes (**Fig. 4B and C**). In contrast, HA-DGKζ^WT^ expression was sufficient to promote extensive blebbing in myotubes (**Fig. 4B and C**). DGKζ^M1^ and the catalytically inactive DGKζ mutant (DGKζ^ΔATP^) induced blebbing to approximately the same extent as DGKζ^WT^ (**Fig. 4A-C**). These results are consistent with RhoA activation being independent of DGKζ kinase activity, as we previously reported (9). To test the effect of the DGKζ C-terminal PDZ-binding motif, we used the DGKζ^FLAG^ mutant, which has an appended FLAG epitope tag that prevents interaction of the motif with syntrophin PDZ domains (**Fig. 4A**) (20). Despite being expressed at levels equivalent to the other DGKζ constructs, DGKζ^FLAG^ did not induce blebbing in myotubes above the level induced by GFP (**Fig. 4C**). These data show DGKζ expression in myotubes, which have low endogenous Rac1 activity, induces substantial membrane blebbing that depends on the DGKζ-syntrophin interaction. Finally, to verify that blebbing resulting from DGKζ expression involves the activation of canonical RhoA signaling, we treated DGKζ-expressing myotubes with Y-27632, an inhibitor specific to the RhoA effector ROCK. DGKζ^M1^-induced blebbing in myotubes was completely blocked by treatment with 10 μM Y-27632, demonstrating DGKζ activates the RhoA-ROCK signaling pathway upstream of membrane blebbing (**Fig. 5**).

**Figure 4.**
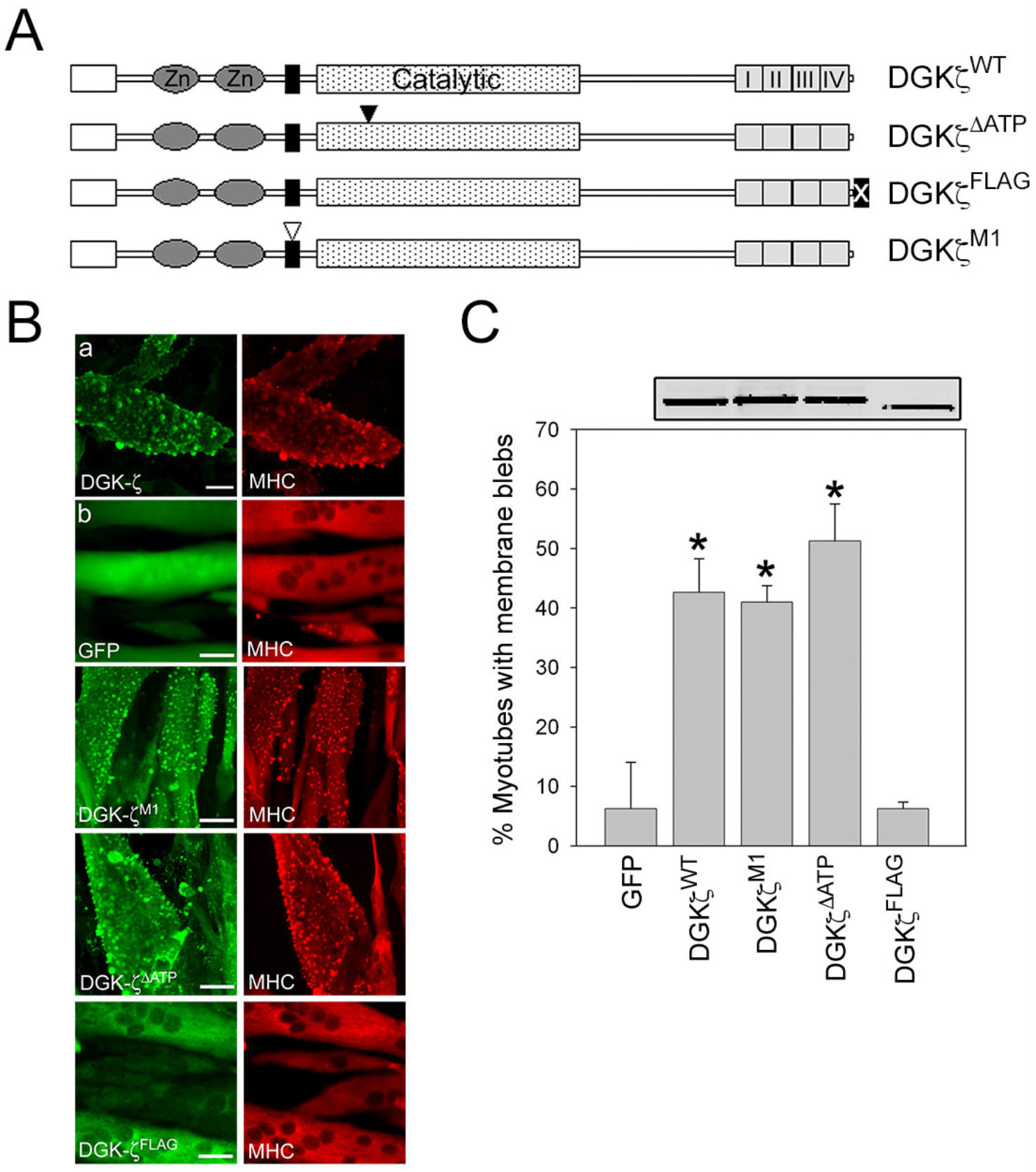
DGKζ expression induces membrane blebbing in C2 myotubes. (**A**) Schematic showing the DGKζ mutants tested. The two zinc fingers (Zn) (dark grey ellipses), MARCKS (black rectangle), catalytic (stippled rectangle) and four ankyrin (light grey rectangles) domains are shown. The open inverted triangle represents four Ser to Asp mutations within the MARCKS domain, which mimics phosphorylation at these sites (DGKζ^M1^). The filled inverted triangle indicates a Gly to Asp mutation in the catalytic domain that completely eliminates DGKζ activity (DGKζ^ΔATP^). The filled box containing an X represents a C-terminal FLAG epitope tag that blocks the interaction with syntrophins (DGKζ^FLAG^). (**B**) Cultures of C2 myotubes differentiated for 48 h were infected with adenoviral vectors harboring the indicated DGKζ constructs or GFP. The cells were fixed 24 h post-infection and were processed for double immunofluorescence microscopy using anti-HA or anti-FLAG and anti-MHC antibodies. Scale bars = 20 μm. (**C**) Graph showing the percentage of myotubes with membrane blebs. Values are the average of at least 3 independent experiments. Error bars indicate S.D. An asterisk indicates a statistically significant difference from GFP control (p < 0.05, two tailed t test). Above the graph is an immunoblot showing the relative protein level of each DGKζ construct as determined using an anti-DGKζ antibody.

**Figure 5.**
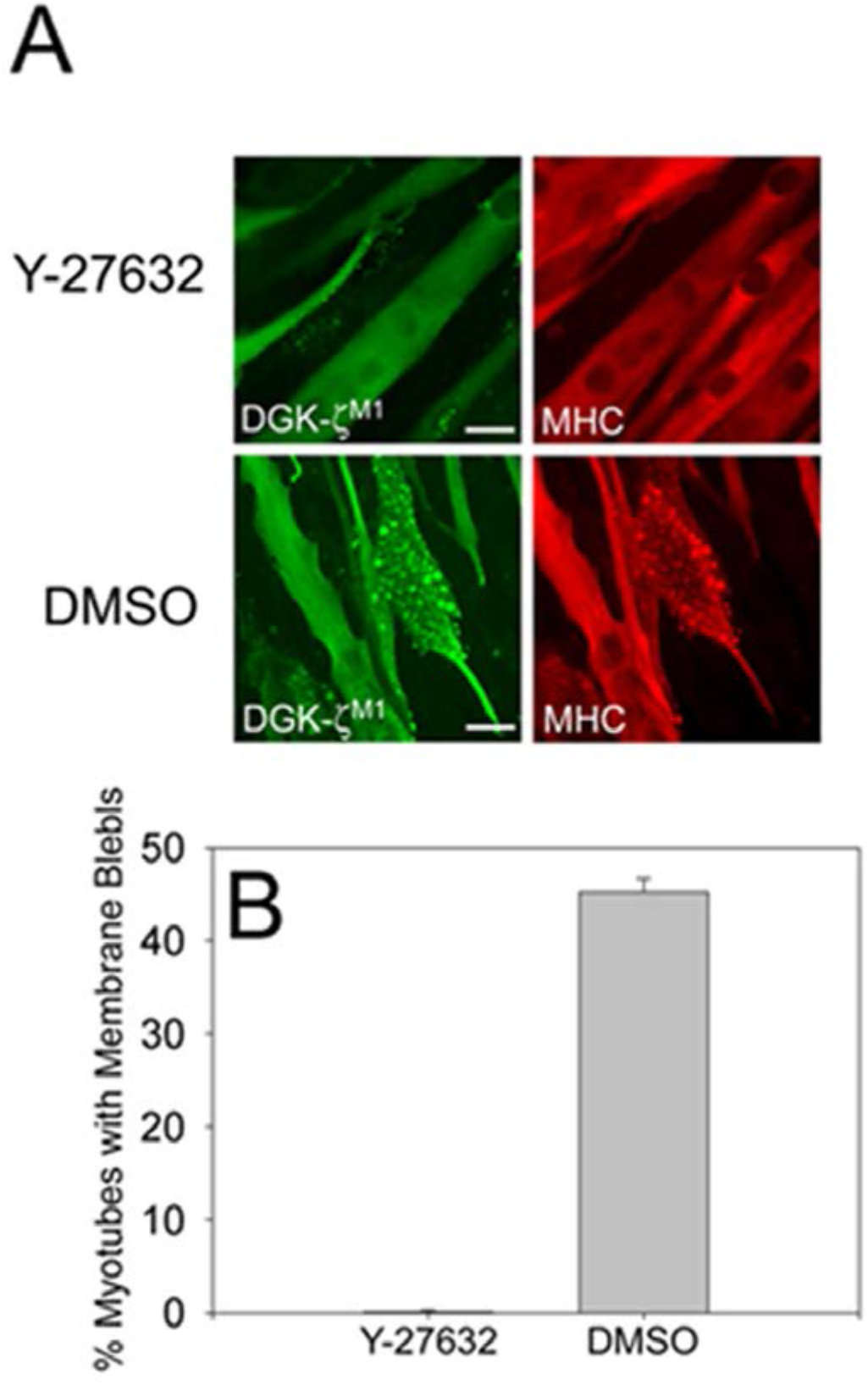
DGKζ-induced membrane blebbing is dependent on ROCK activity. (**A**) Representative images of C2 myotubes expressing HA-DGKζ^M1^ and treated with either 10 μM Y-27632 or DMSO. Cells were fixed 24 hours post-infection and were double labeled with anti-HA and myosin heavy chain (MHC). Scale bars = 20 μm. (**B**) Graph showing the percentage of infected, blebbing myotubes in each condition. Values are the average of three independent experiments. Error bars represent S.E.M. The treated condition was statistically significant different from DMSO (*p* < 0.05) by one-tailed *t* test.

### DGKζ-Induced RhoA Activation Requires C-terminal PDZ Interactions

To investigate the mechanistic basis for the failure of DGKζ^FLAG^ to induce blebbing in myotubes, we tested its ability to rescue RhoA activity in DGKζ-null MEFs, which have reduced levels (~50%) of active RhoA compared to wild type MEFs (9). We assayed the level of GTP-bound (active) RhoA in lysates of uninfected null cells or null cells infected with adenovirus harboring HA-tagged DGKζ^WT^, DGKζ^ΔATP^ or DGKζ^FLAG^ using an effector pull-down assay (26) (**Fig. 6**). RhoA activity was increased ~2-fold in lysates of null cells infected with either DGKζ^WT^ or DGKζ^ΔATP^ but RhoA activity in DGKζ^FLAG^-expressing cells was not significantly different from uninfected DGKζ-null cells. In the same experiment, DGKζ^WT^ and DGKζ^FLAG^ increased Rac1 activity by ~2-fold, whereas DGKζ^ΔATP^ failed to rescue Rac1 activity, consistent with our previously published findings (9). These results suggest a functional PDZ binding motif is required for DGKζ-induced RhoA activation but is dispensable for Rac1 activation.

**Figure 6.**
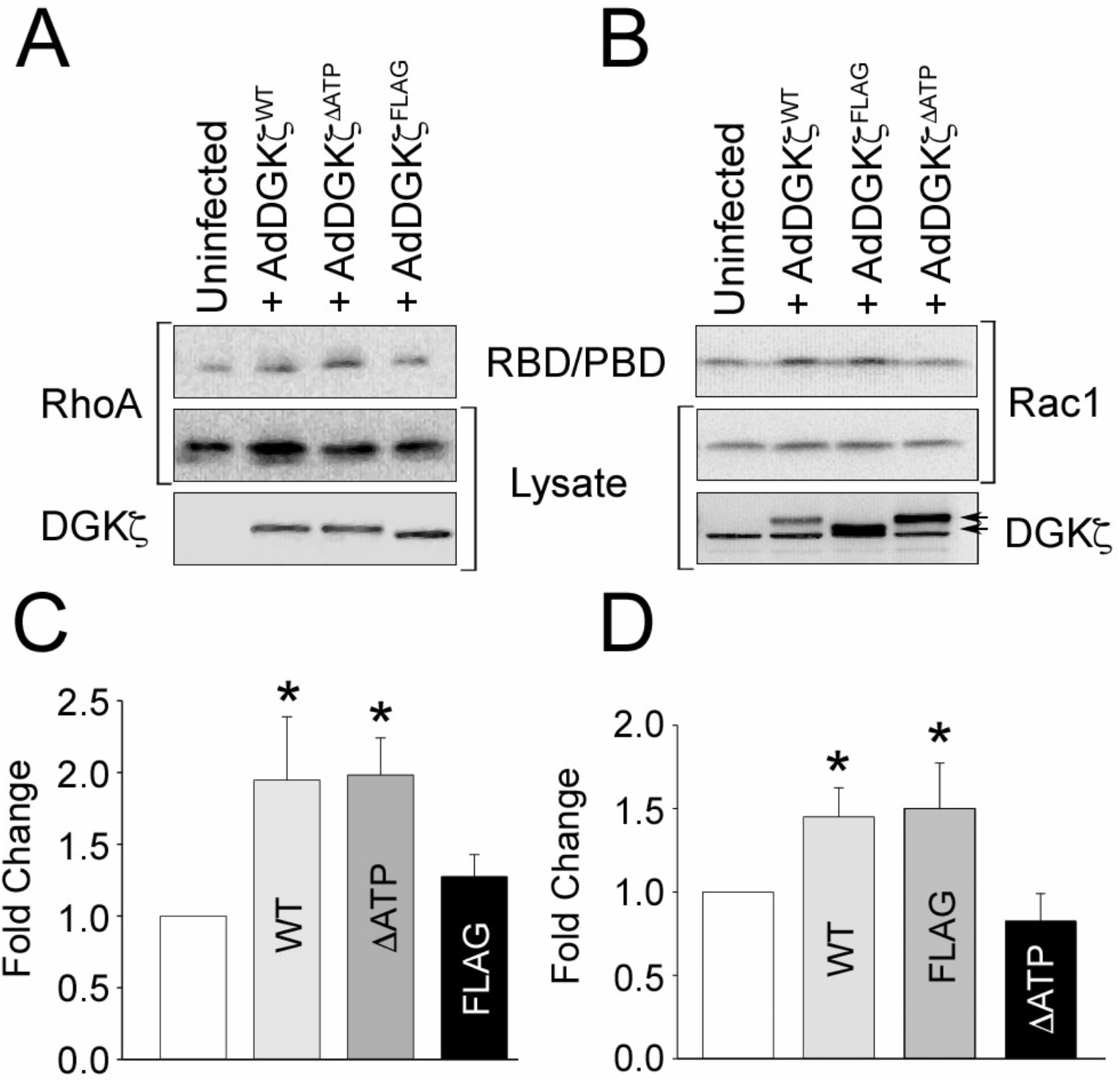
Rescue of RhoA activation requires a functional C-terminal PDZ binding motif. DGKζ-null MEFs were infected with adenovirus bearing wild type HA-DGKζ (AdDGKζ^wt^), a catalytically inactive (kinase dead) mutant (AdDGKζ^kd^), or a mutant with a C-terminal FLAG epitope tag that blocks interaction with syntrophins (AdDGKζ^FLAG^). (**A**) Serum-stimulated cells were lysed and the extracts incubated with an immobilized GST fusion protein of the Rho binding domain of Rhotekin (GST-RBD) to capture GTP-bound RhoA. Bound proteins were analyzed by immunoblotting for RhoA. (**B**) The cells were stimulated for 5 min with 50 ηg/ml PDGF. Global Rac1 GTPase activity was assayed by pulldown with a GST fusion protein of p21-binding domain (PBD) of PAK1, followed by immunoblotting for Rac1. In A and B, the cell lysates were immunoblotted for DGKζ and RhoA or Rac1. In **B**, arrows indicate the different sized, exogenously expressed DGKζ proteins. The lower band is non specific. (**C and D**) Graphs showing the quantification of active RhoA and Rac1, under the respective immunoblots, by densitometric analysis. Values are the average fold change ± S.E.M. from three independent experiments. The asterisks indicate a statistically significant difference from uninfected null cells (*p* < 0.05) by one-tailed *t* test.

## Discussion

The Rho GTPases Rac1 and RhoA are sequestered in separate signaling complexes bound to their common inhibitor RhoGDI, which maintains them in their inactive state. Their selective dissociation from RhoGDI allows the precise control of downstream responses like changes in actin organization in response to extracellular signals. DGKζ is an integral part of both dissociation mechanisms, but has somewhat different roles in each complex; its catalytic activity is required for Rac1 dissociation but is dispensable for RhoA dissociation, and instead, DGKζ functions as a scaffold (8,9)}. Nevertheless, DGKζ lies upstream of both Rac1 and RhoA activation and is therefore in a key position to regulate the balance of their respective signaling pathways. The main finding of this work is that DGKζ achieves this regulation in part by PKCα-mediated phosphorylation of the DGKζ MARCKS domain, which functions as an intramolecular switch that promotes the interaction of DGKζ with RhoA and simultaneously decreases its interaction with Rac1. Since MARCKS phosphorylation also attenuates DGKζ catalytic activity (18) and DGKζ catalytic activity is required to activate Rac1 but not RhoA, decreasing its activity would lead to a reduction in Rac1 activity and a relative increase in RhoA activity. Moreover, since active Rac1 directly and indirectly inhibits RhoA activity by several different mechanisms (27), this effect would be amplified by decreased inhibition of RhoA by Rac1. Indeed, under conditions of reduced Rac1 activity, RhoA activation becomes the default pathway. This potentially explains why DGKζ^WT^ and DGKζ^ΔATP^ were as effective as DGKζ^M1^ at inducing membrane blebbing in C2 myotubes, which have low endogenous Rac1 levels. Figure 7A summarizes our working model for how MARCKS domain phosphorylation differentially regulates Rac1 and RhoA signaling.

**Figure 7.**
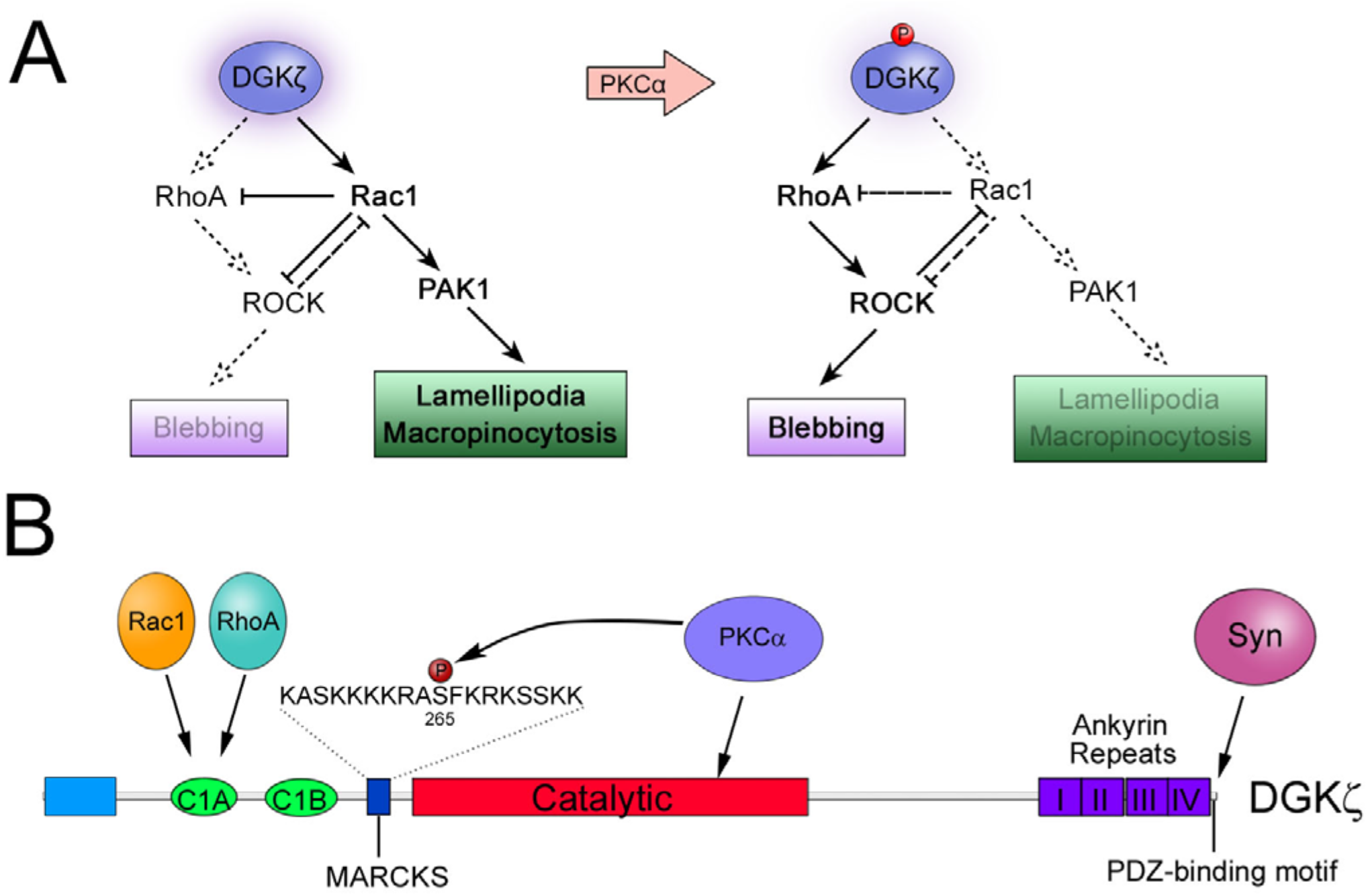
**(A) Proposed Switch–like Mechanism to Regulate the Balance of Rac1 and RhoA Signaling.** We showed previously that DGKζ catalytic activity is required for Rac1, but not RhoA, activation. (**Left**) High DGKζ activity (*purple glow*) favors Rac1 activation and leads to inhibition of Rho signaling by Rac1. (**Right**) PKCα-mediated phosphorylation of the MARCKS domain decreases DGKζ catalytic activity, which decreases Rac1 activation (*dashed arrow*), leading to decreased inhibition and a relative increase in RhoA signaling. (**B**) **Schematic diagram of DGKζ** showing the N-terminal domain (blue), C1 domains (green), MARCKS domain (dark blue), catalytic domain (red), ankyrin repeats (purple) and the C-terminal PDZ-binding motif. The binding sites for Rac1, RhoA, PKCα and syntrophin (Syn) are indicated. PKCα binds to a region in the catalytic domain and phosphorylates DGKζ at Ser-265 in the MARCKS domain. Both Rac1 and RhoA bind to the C1A domain and their binding is differentially regulated by MARCKS phosphorylation.

PKC activity has previously been implicated in membrane blebbing in pancreatic acinar cells (28) and in muscarinic agonist-induced spectrin redistribution accompanied by bleb formation (29). These studies were limited by the use of the broad-spectrum kinase inhibitor staurosporine, so in retrospect a role in blebbing cannot be definitively attributed to PKCα. Nevertheless, other studies support the idea that PKCα is a component of the blebbing machinery. The MARCKS protein, from which the domain in DGKζ gets its name, is a PKCα substrate that cycles on and off membranes by a mechanism termed the myristoyl–electrostatic switch (30). At the plasma membrane, MARCKS binds to and sequesters acidic phospholipids including PI(4,5)P_2_. A mutant MARCKS protein, in which the electrostatic switch was replaced by a constitutive membrane targeting sequence, generated dynamic membrane blebs when expressed in cells, implicating MARCKS and PKCα in the blebbing response (31). Our findings firmly connect PKCα activity to the activation of membrane blebbing and indicate that the PKCα and RhoA pathways intersect at the level of DGKζ regulation of RhoA activity. In our studies, treatment of C2 myotubes with PMA failed to activate membrane blebbing (unpublished observations) suggesting other upstream signals are required to initiate this response.

### Role of the DGKζ C-terminal PDZ-binding Motif

Not only did MARCKS domain phosphorylation increase the interaction with RhoA but it also increased binding to α1-syntrophin, an effect mediated by the DGKζ C-terminal PDZ-binding motif and the α1-syntrophin PDZ domain. At present, the biological significance of this increased interaction is uncertain, but it is consistent with the idea that a DGKζ-RhoA complex includes syntrophin. Indeed, the DGKζ C-terminal PDZ-binding motif was required to rescue RhoA activity in DGKζ-null cells, suggesting syntrophin interaction is required for optimal RhoA activation. One possibility is that syntrophin regulates the subcellular localization of a DGKζ/RhoA complex. We showed previously that syntrophins regulate DGKζ subcellular localization in several different mammalian cell types (17,19,20). In skeletal muscle cells, co-expression of α1-syntrophin potentiated the plasma membrane localization of DGKζ^M1^ suggesting the two are coordinately regulated by MARCKS phosphorylation. This would serve as an effective mechanism to transport DGKζ/RhoA complex to the plasma membrane and bring it into proximity to GEFs and membrane bound effectors.

In a previous study, we showed that expression of a catalytically inactive DGKζ mutant in DGKζ-null MEFs could restore active RhoA to near wild type levels (9). Although the DGKζ^FLAG^ mutant was unable to rescue RhoA activity in DGKζ-null MEFs, it was able to rescue Rac1 activity, indicating the C-terminal PDZ interaction is not required for Rac1 activation. This functional requirement for syntrophin interaction further differentiates the two signaling complexes.

### MARCKS Domain Phosphorylation: A Multifunctional Switch

The differential binding to Rac1 and RhoA adds to a growing number of DGKζ interactions affected by MARCKS domain phosphorylation. Both Rac1 and RhoA bind to the C1 domain (C1A) of DGKζ (8,9), which is located close to the N-terminus (**Fig. 7B**). The MARCKS domain is situated just downstream of a second C1 domain (C1B), so it is perhaps not surprising that phosphorylation affects binding to nearby domains. Remarkably, MARCKS domain phosphorylation also affects syntrophin binding to the PDZ-binding motif at the extreme C-terminus, suggesting significant three-dimensional structural changes accompany PKCα-mediated phosphorylation. Consistent with this idea, PKCα binding to the latter half of the DGKζ catalytic domain is abolished by MARCKS phosphorylation, relieving the inhibition of PKCα imposed by DGKζ and allowing prolonged PKCα activation (13). MARCKS phosphorylation also attenuates DGKζ activity by approximately 50%, which limits its ability to metabolize signaling DAG. More importantly for Rac1 activation, this decreases PA production required for stimulating PAK1-mediated release of Rac1 from RhoGDI (8). Finally, there are likely yet-to-be identified signaling pathways that stimulate phosphatase activity to reverse MARCKS domain phosphorylation, thereby switching the preference of DGKζ back from RhoA to Rac1 signaling. This Rho GTPase signaling network has been shown mathematically and experimentally to exhibit a bi-stable response to perturbations (22,32). Thus, the level of MARCKS domain phosphorylation could function as a rheostat to tune Rac1/RhoA signaling and change the signaling output of the Rho GTPase network.

The balance between Rac1 and RhoA signaling underpins two different modes of cell migration. Rac1 signaling promotes a mesenchymal mode characterized by an elongated shape that requires extracellular proteolysis at cellular protrusions. In contrast, RhoA signaling drives an amoeboid mode in which movement is independent of proteases, cells have a rounded morphology with no obvious polarity, and the plasma membrane undergoes active blebbing driven by actomyosin contractility (33,34). Inhibitory signals suppress the activity of the opposing pathway so that one pathway predominates; however, cells are able to switch between these two modes of movement. Our findings demonstrate DGKζ occupies a central node at the apex of the Rac1 and RhoA signaling pathways and that PKCα-mediated phosphorylation of DGKζ is a switch that favours RhoA over Rac1 signaling. Thus, the phosphorylation of DGKζ by PKCα may be one of the stimuli that triggers the interconversion between migratory modes.

## Experimental Procedures

### Antibodies

Monoclonal and polyclonal anti-HA, and monoclonal anti-tubulin and anti-actin antibodies were purchased from Sigma-Aldrich (St. Louis, MO). Monoclonal anti–c-myc antibody was from Roche Applied Science (Indianapolis, IN). An affinity-purified polyclonal antibody raised against the N-terminus of DGKζ has been described previously (15). Anti-GFP and anti-myosin heavy chain (MHC) antibodies were from Santa Cruz Biotechnology, Inc. Alexa Fluor 488– and 594–conjugated secondary antibodies and phalloidin were purchased from Invitrogen (Carlsbad, CA). HRP-conjugated anti-rabbit and anti-mouse secondary antibodies were from Jackson ImmunoResearch Laboratories (West Grove, PA). The Rac1 monoclonal antibody 102 was purchased from BD Transduction Laboratories and the RhoA monoclonal antibody (26C4) was from Santa Cruz Biotechnology, Inc. Monoclonal antibody 1351 raised against syntrophin was a gift from Dr. Stanley Froehner (University of Washington, Seattle, WA).

### Plasmids

The Rhotekin Rho binding domain (RBD) construct was described previously (Ren et al., 1999 blue right-pointing triangle). Plasmids encoding wild-type DGKζ and a catalytically inactive mutant (DGKζ^ΔATP^), both with three tandem, N-terminal HA epitope tags, as well as DGKζ with a C-terminal FLAG epitope tag (DGKζ^FLAG^), have been described previously (15,20). Rac1^V12^ and Rac1^N17^ were cloned into pGEX-4T-1 as described previously (8,19). RhoA^V14^ and RhoA^N19^ were obtained from the Missouri S&T cDNA Resource Center (www.cdna.org) and were subcloned into pGEX-4T-3 as described (9). The PDZ domain of α1-syntrophin was cloned into pGEX-4T-1 as described previously (20).

### Cell culture

Immortalized wild type and DGKζ-null mouse embryonic fibroblast (MEF) lines have been described previously (8). MEFs were cultured at 37°C in 5% CO_2_ in DMEM high-glucose supplemented with 10% fetal bovine serum (FBS), 2 mM L-glutamine, 100 U/ml penicillin, and 100 U/ml streptomycin. C2C12 myoblasts were plated on dishes coated with Matrigel (Collaborative Research, Bedford, MA) or collagen, as indicated, and culture at 37°C in 5% CO_2_ in DMEM high-glucose supplemented with 10% FBS, 100 U/ml penicillin-streptomycin, and 2 mM L-glutamine. To induce differentiation into myotube fibers, C2C12 plates were grown to 80-100% confluence and then switched to medium containing 5% horse serum. For PKC activation, cells were serum starved overnight and stimulated with 100 ηM with phorbol-12-myristate-13-acetate (PMA) or vehicle (dimethyl sulfoxide) for 10 min. In some experiments, cells were pretreated 30 min with 1 μM Gö6976, a potent PKCαβ-specific inhibitor. For Rac1 and ROCK inhibition experiments, cells were treated with 100 μM NSC 23766 or 10 μM Y-27632, respectively.

### Transfection and adenoviral infection

C2 myoblasts were transfected 18-24 h after plating at 70-80% confluence by using FuGENE 6 (Roche Diagnostics, Indianapolis, IN) according to the manufacturer’s instructions. For transfections of myoblasts plated on glass coverslips, 1 μg of purified DNA was added to 3 μl of FuGENE 6 (3:1 ratio) diluted in serum-free DMEM to a final volume of 100 μl. The mixture was incubated for 20 min at room temperature and added to 3 ml of growth medium in 35-mm dishes containing the coverslips. MEFs were plated onto collagen-coated plates and transfected at 60-80% confluency using FuGENE 6 (Roche, Indianapolis, IN) according to manufacturer’s instructions. The cloning and production of adenoviral constructs have been described previously (19). For adenoviral overexpression experiments, fibroblasts, myoblasts and differentiated myotubes were infected at a multiplicity of infection of 100 for 1 h at 37°C. Cells were incubated for an additional 24–36 h under standard growth conditions.

### Immunofluorescence microscopy

Briefly, cells were rinsed with PBS (pH 7.4) before fixation with 4% paraformaldehyde for 15 min at 37°C. After cell permeabilization using 0.5% Triton X-100 in PBS for 10 min, cells were incubated in blocking buffer (filtered 1% BSA in PBS) for 30 min at room temperature. Primary antibodies were diluted in blocking buffer at a ratio of 1:100, unless otherwise indicated. Secondary fluorescent conjugated antibodies were diluted 1:300. F-actin fibers were stained using Alexa Fluor 488- or 594–conjugated phalloidin, which was diluted in blocking buffer at a ratio of 1:1000. Coverslips were mounted onto glass slides using Fluoromount-G (Southern Biotech) and sealed with nail polish. Images were obtained using a charge-coupled device (CCD) camera on an Axioskop 2 microscope with AxioVision software (Carl Zeiss, Jena, Germany).

### Quantification of membrane blebbing

Blebbing was quantified by counting the number of cells with and without membrane blebs in 20 randomly selected fields per coverslip, viewed with a 20X objective. At least two coverslips were counted per condition, with at least 150 cells counted per experiment. A cell undergoing blebbing was defined as the cell having at least one third of the surface covered in blebs.

### RhoA/Rac1 activity assays and GST pulldown assays

The expression and purification of GST fusion proteins was performed as described previously (8,9,20). The level of GTP-bound Rac1 or RhoA was measured by pulldown assay with a GST fusion protein of the p21 binding domain (PBD) of PAK1 or the Rho binding domain (RBD) of Rhotekin, respectively, as we have done before (8,9). Cells were serum-starved overnight and then stimulated with serum for 10 min or with 50 ng ml^−1^ platelet derived growth factor (PDGF) for 5 min to activate RhoA and Rac1, respectively. The medium was quickly removed, and the cells were immediately harvested in chilled lysis buffer (50 mM Tris-HCl, pH 7.4, 150 mM NaCl, 1% Triton X-100, 50 mM MgCl_2_, and protease inhibitors). Cell lysates were centrifuged at 18,000 × g for 15 min at 4°C. Equivalent amounts of protein were incubated with GST-RBD or GST-PBD beads for 30 min at 4°C. For GST pulldown assays, cells expressing HA-tagged DGKζ constructs were lysed, and equal amounts of protein were incubated with immobilized GST-fusion proteins for 1-2h at 4°C and washed a minimum of four times with ice cold lysis buffer. The beads were collected, washed several times with lysis buffer and then boiled in reducing Laemmli sample buffer. The eluted proteins were analyzed by SDS-PAGE and immunoblotting.

### Immunoprecipitation

Immunoprecipitations were carried out essentially as described previously (20). Briefly, cells were lysed in 50 mM Tris-HCl, pH 7.5, 50 mM NaCl, 5 mM MgCl2, 1% NP-40, 5% glycerol, 1 mM dithiothreitol (DTT), and protease inhibitors and centrifuged at 18,000 x g for 10 min at 4°C. Equivalent amounts of protein (~1 mg) were incubated with 5 μg antibody or control rabbit immunoglobulin G (IgG) for 4h at 4°C. Then, 40 μl of 50% protein G agarose slurry was added for 1.5 h. The beads were washed with lysis buffer, resuspended in reducing Laemmli buffer, and analyzed by SDS-PAGE and immunoblotting.

### Western blotting and quantification of digital images

Proteins were separated by SDS-PAGE and transferred to PGDF membranes as described (Ard et al. 2012). Blots were stained with Ponceau S to record the total protein loaded in each lane before proceeding with antibody incubations and chemiluminescent detection. Antibodies were diluted in 5% skim milk powder dissolved in 25 mM Tris pH 7.5, 150 mM NaCl and 0.1% Tween-20. Images were captured with a LI-COR Odyssey digital imaging system (LI-COR Biosciences, Inc.) and the intensity of the bands was analyzed using Image Studio software. The raw data was imported into Excel, analyzed and then exported to SigmaPlot 12 for graphing and statistical analysis.

## Data Availability

All data are contained within the manuscript.

## Acknowledgment

We are grateful to Helga Agah for excellent technical assistance.

## Abbreviations

CBB: Coomassie Brilliant Blue
DAG: diacylglycerol
DGKζ: diacylglycerol kinase ζ
GDI: guanine nucleotide dissociation inhibitor
GST: glutathione S-transferase
HA: hemagglutinin
MARCKS: myristoylated alanine-rich C kinase substrate
MEF: mouse embryonic fibroblast
MHC: myosin heavy chain
PA: phosphatidic acid
PBD: p21 binding domain
PKCα: protein kinase Cα
RBD: Rho binding domain
RhoGDI: Rho guanine nucleotide dissociation inhibitor
SD: standard deviation
SEM: standard error of the mean

